## Supplemental Figures for "DIPPER: a spatiotemporal proteomics atlas of human intervertebral discs for exploring ageing and degeneration dynamics"

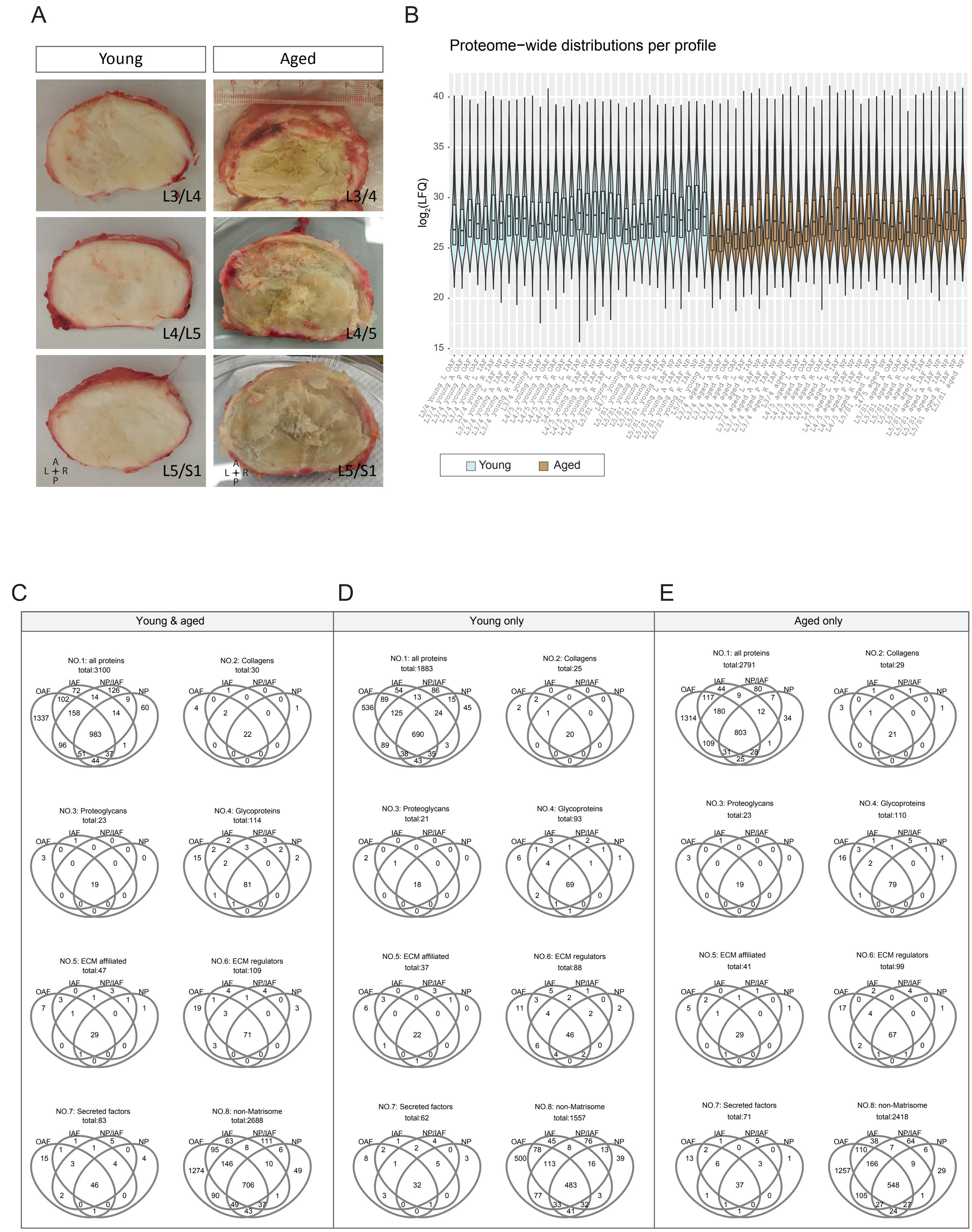


**Figure S1.** (A) Gross images of the young and aged cadaveric discs. (B) Proteome-wide distributions per profile across all 66 profiles. The profiles were named with levels, ages, directions, and compartments. (C)-(E) Venn diagrams of detected proteins among the four major IVD compartments (OAF, IAF, NP/IAF, and NP), per age-group, and per protein category.

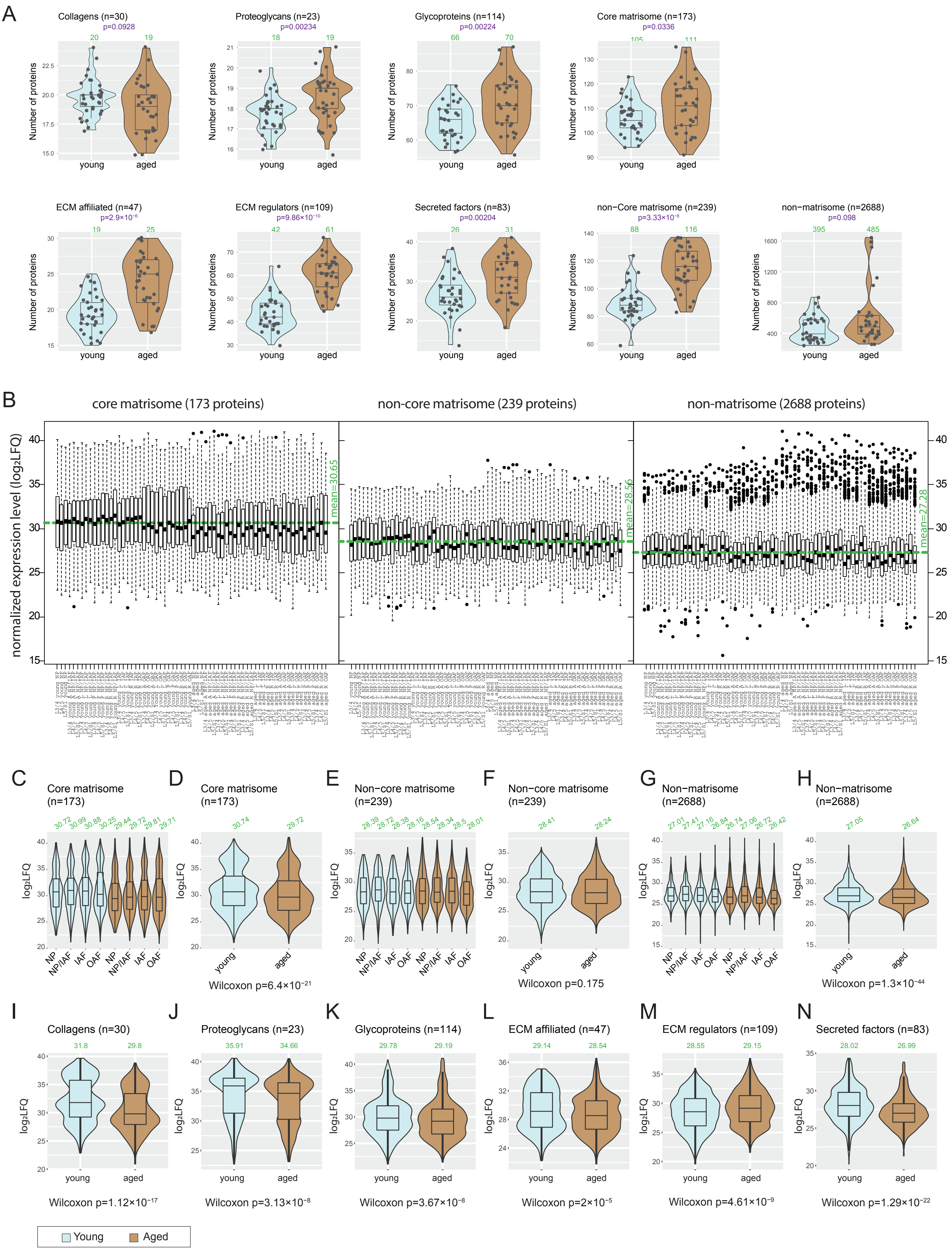


**Figure S2.** (A) Violin-plots showing numbers of proteins detected per age-group, for all categories of ECM and non-ECM proteins. The green numbers on top of each violin show the median number of proteins detected per respective sample group. (B) Box-plots showing the expression levels of core-matrisome, non-core matrisome, and non-matrisome proteins. Horizontal green line indicates average. (C)-(H) Violin plots showing the expression levels of major ECM categories across compartments and age-groups. (I)-(N) Violin plots showing the expression levels of sub-categories of ECM proteins across age-groups. The green numbers on top of each violin show the median number of proteins detected per respective sample group.

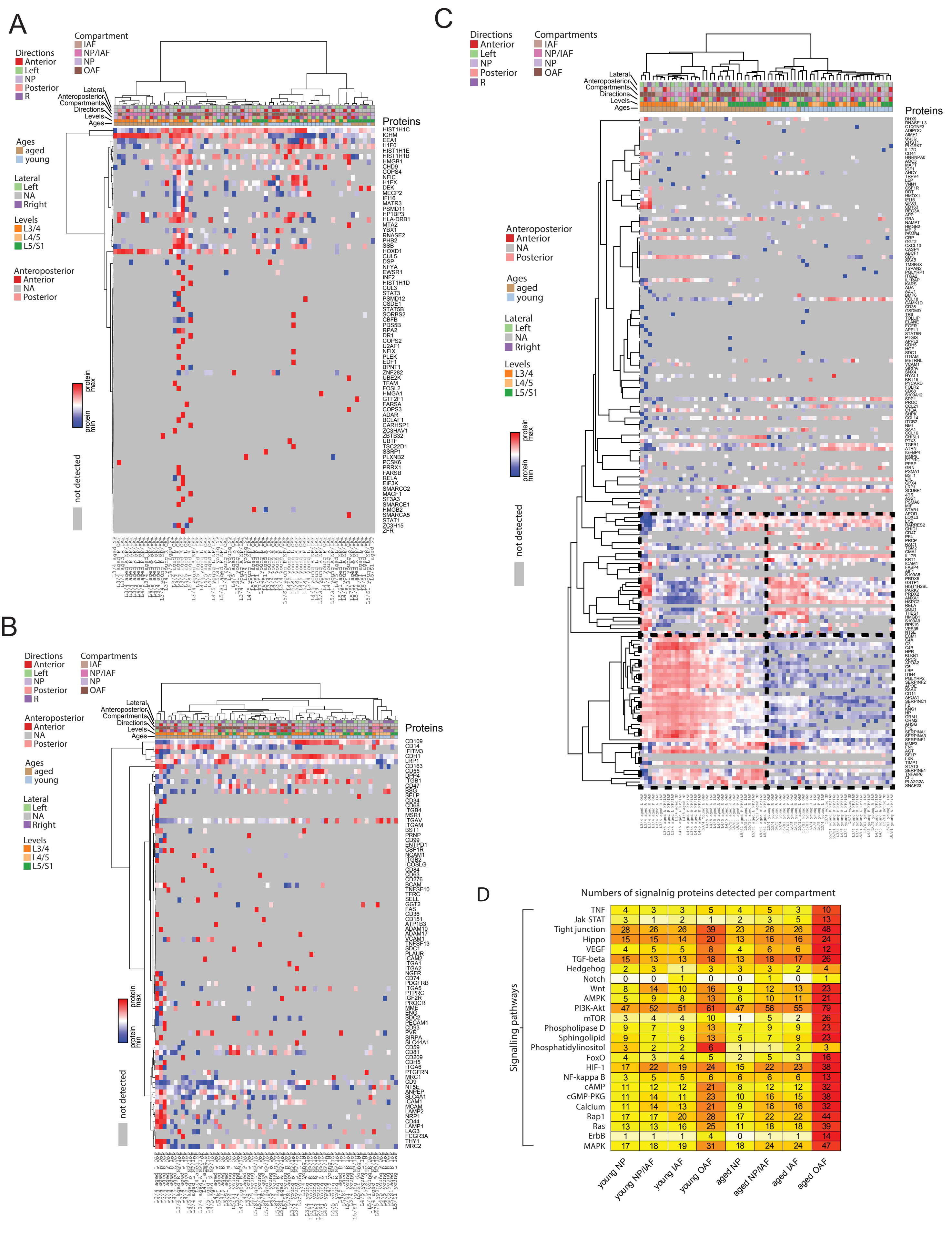


| | young NP | young NP/IAF | young IAF | young OAF | aged NP | aged NP/IAF | aged IAF | aged OAF |
|---|---|---|---|---|---|---|---|---|
| TNF | 4 | 3 | 3 | 5 | 4 | 5 | 3 | 10 |
| Jak-STAT | 3 | 1 | 2 | 1 | 2 | 3 | 5 | 13 |
| Tight junction | 28 | 26 | 26 | 39 | 23 | 26 | 26 | 48 |
| Hippo | 15 | 15 | 14 | 20 | 13 | 16 | 16 | 24 |
| VEGF | 4 | 5 | 5 | 8 | 4 | 6 | 6 | 12 |
| TGF-beta | 15 | 13 | 13 | 18 | 13 | 18 | 17 | 26 |
| Hedgehog | 2 | 3 | 1 | 3 | 3 | 3 | 2 | 4 |
| Notch | 0 | 0 | 1 | 0 | 0 | 1 | 0 | 1 |
| Wnt | 8 | 14 | 10 | 16 | 9 | 12 | 13 | 23 |
| AMPK | 5 | 9 | 8 | 13 | 6 | 10 | 11 | 21 |
| PI3K-Akt | 47 | 52 | 51 | 61 | 47 | 56 | 55 | 79 |
| mTOR | 3 | 4 | 4 | 10 | 1 | 5 | 2 | 26 |
| Phospholipase D | 9 | 7 | 9 | 13 | 7 | 9 | 9 | 23 |
| Sphingolipid | 9 | 7 | 6 | 13 | 5 | 7 | 9 | 23 |
| Phosphatidylinositol | 3 | 2 | 2 | 6 | 1 | 1 | 2 | 3 |
| FoxO | 4 | 3 | 4 | 5 | 2 | 5 | 3 | 16 |
| HIF-1 | 17 | 22 | 19 | 24 | 15 | 22 | 23 | 38 |
| NF-kappa B | 3 | 5 | 5 | 6 | 6 | 6 | 6 | 13 |
| cAMP | 11 | 12 | 12 | 21 | 8 | 12 | 12 | 32 |
| cGMP-PKG | 11 | 14 | 11 | 23 | 10 | 16 | 15 | 38 |
| Calcium | 11 | 14 | 13 | 21 | 9 | 16 | 16 | 32 |
| Rap1 | 17 | 17 | 20 | 28 | 17 | 22 | 22 | 44 |
| Ras | 13 | 13 | 16 | 25 | 11 | 18 | 18 | 39 |
| ErbB | 1 | 1 | 1 | 4 | 0 | 1 | 1 | 14 |
| MAPK | 17 | 18 | 19 | 31 | 18 | 24 | 24 | 47 |

**Figure S3.** Profiles and their hierarchical clustering patterns of all detected transcription factors or DNA-binding proteins (A), cell surface markers (B), inflammatory proteins (C), and numbers of signalling proteins detected per compartment and age-group (D) in the data. In (D), the color scale corresponds to proteins in overlap within each entry divided by total number of proteins in the pathway.

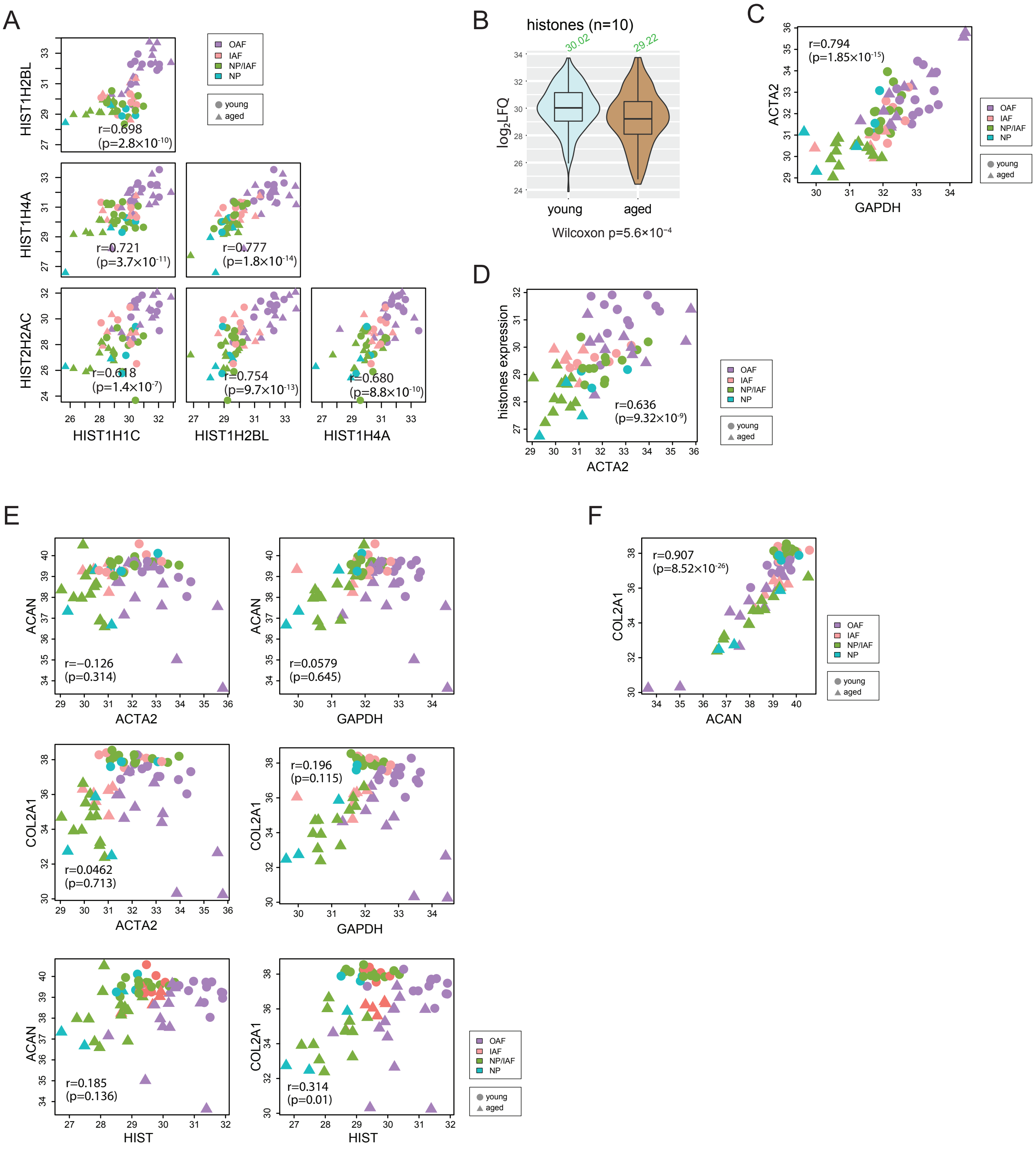

**Figure S4.** Histones and housekeeping genes reflect cellularities. (A) Scatter-plots showing the co-expression of four histone proteins that were detected in over 60 profiles. (B) Violin plot showing the expression levels of the histones across age-groups. (C) Scatter plot showing the co-expression between ACTA2 and the average of histones. (D) Scatter plot showing the co-expression between ACTA2 and GAPDH. (E) Scatter plots showing the co-expression between ACTA2, GAPDH, and histones, and COL2A1, and ACAN. (F) Scatter plot showing the co-expression between COL2A1 and ACAN. All values are in $log_2(LFQ)$. r is Pearson correlation coefficient.

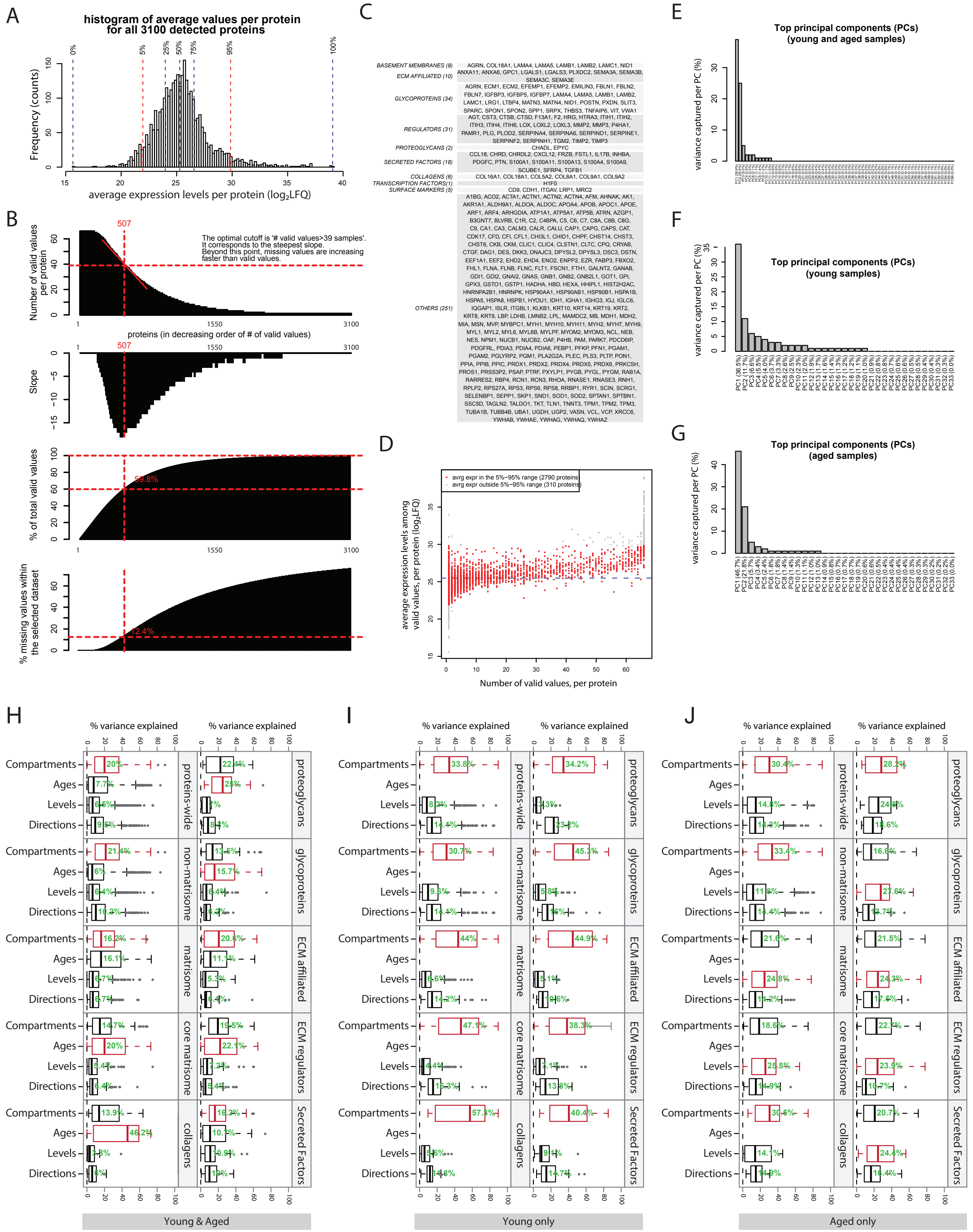


**Figure S5.** (A) Histogram and key percentiles of the average expression levels of all 3100 detected proteins. Average was calculated based on the valid samples of each protein. (B) Selecting an optimal cutoff above which proteins will be used for performing PCA. Upper panel, proteins were ordered in decreasing order by its number of valid values. Second panel: the slope of the upper panel. Third panel: the fraction of all valid values in the whole data-set captured per each cutoff. An optimal cutoff corresponding to the cutoff of '>39 valid values' was found where the slope is steepest. Lower panel: the fraction of missing values within the selected dataset at each cutoff. (C) A list of the proteins (categorized by their functional families) that both meet the criteria in (B) and fall in the 5%~95% range in (A). (D) A scatter plot showing the positive relation between number of valid values per protein, and the average expression level per protein. (E)-(G), the percentages of variance captured by the top principal components (PCs) in the whole dataset (E), young samples only (F) and aged samples only (G). (H)-(J) Horizontal box plots showing the percentages of variance explained by four phenotypic factors, for different scopes of protein sets. (H) young and aged combined. (I) Young only. (J) Aged only.

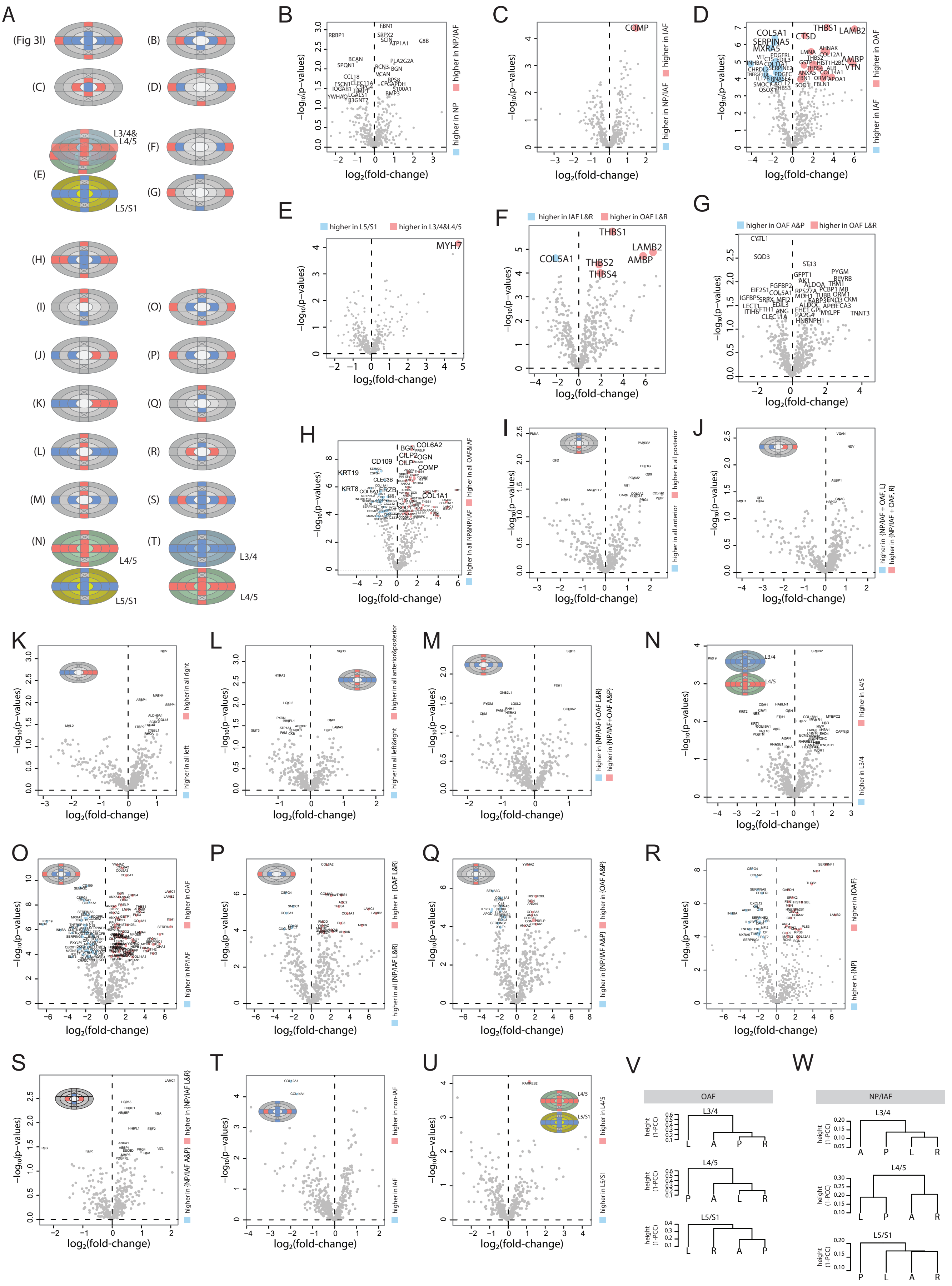

**Figure S6.** Additional comparisons within the young samples. (A) Schematic diagrams showing the comparisons between different groups of samples. (B)-(U), volcano plots showing the differentially expressed proteins for each comparison listed in (A). (V-W) Dendrograms showing the clustering patterns of four samples corresponding to left (L), right (R), anterior (A), and posterior (P) directions, in OAF (V) and NP/IAF (W), respectively.

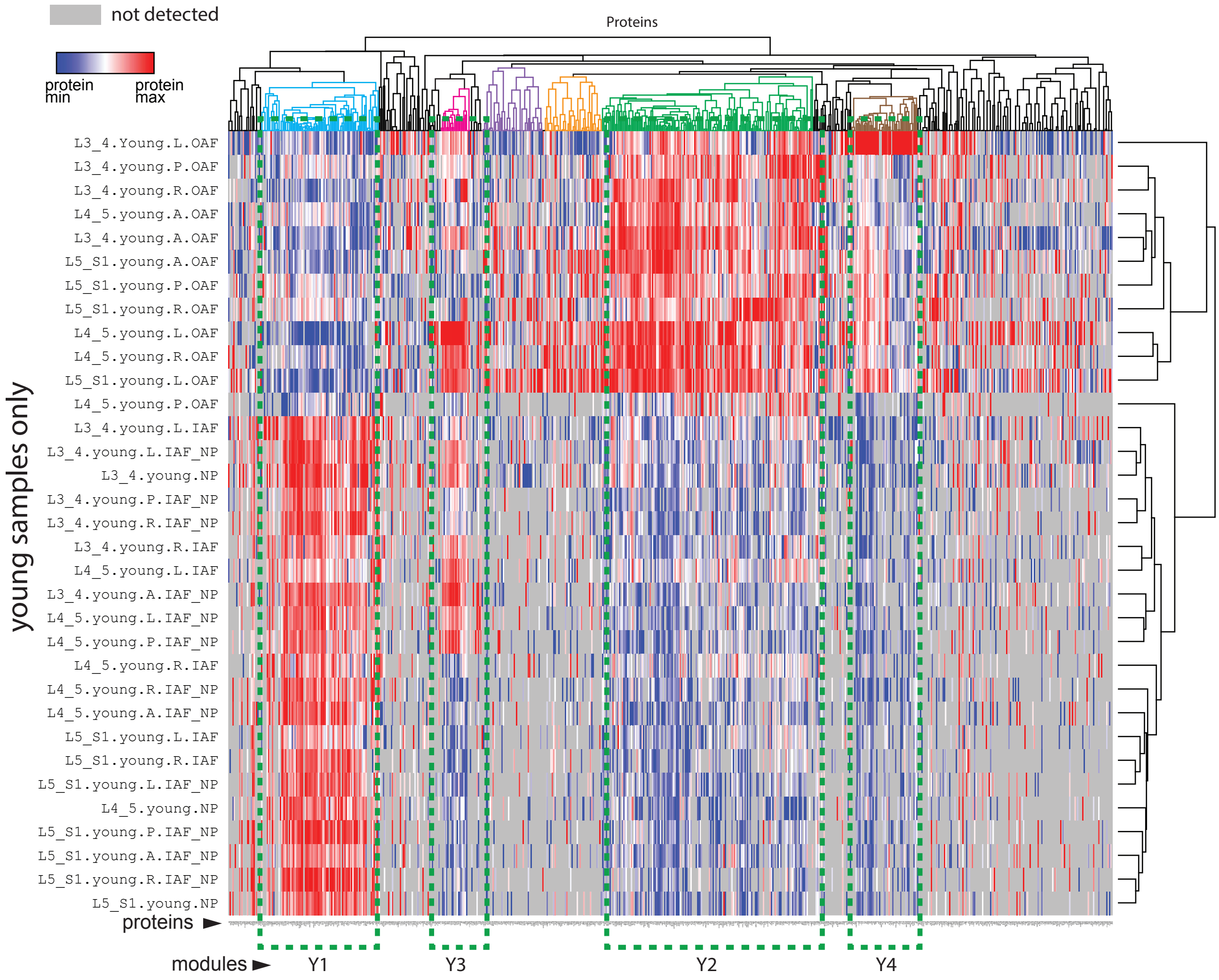


**Figure S7.** A profile-protein bi-clustering heatmap of 671 differentially expressed proteins identified in 20 two-group comparisons within the young samples. For each of the comparisons, a DEP could come from three sources: statistical comparisons, fold-changes, or exclusive expressions in one group only (Methods). Four protein modules were identified: Y1~Y4.

A

**Proteins in Y1 (n=96):**

DAG1, DNAJC3, FIBIN, LPL, NUCB1, CPQ, GARS, PREP, RSU1, CLEC3B, EFEMP1, KRT19, FN1, PRG4, CXCL12, B4GAT1, KAL1, CSPG4, INHBA, SLIT3, **CDH1**, SLPI, CYTL1, IGFBP5, CHADL, ISLR, COL5A2, MATN3, EFHD2, VIT, SMOC2, PCOLCE2, BCAN, CST3, CCDC80, ATRN, EMILIN1, EPHX1, RNASE4, PCOLCE, COL11A2, OAF, PLOD1, XYLT1, DKK3, CLSTN1, EDIL3, MFI2, COL5A1, SERPINA5, C1S, CP, SERPING1, FRZB, APOD, COL11A1, TXNRD1, IL17B, QSOX1, SERPINE2, TNFRSF11B, MXRA5, PDGFRL, **CD109**, KRT8, SEMA3C, TIMP3, CFB, HYAL1, ITIH5, SMOC1, PXYLP1, COL3A1, THBS3, VAPA, GALNT2, CCT4, **MECP2**, GDF6, CHRD, CHRDL2, UBXN10, TIMP1, PDGFC, IGFBP7, CYFIP1, SLC44A2, PPP2R1A, SEPP1, AKR1C1, MGAT1, VEGFA, GBA, RPL6, ENPP2, SCUBE1

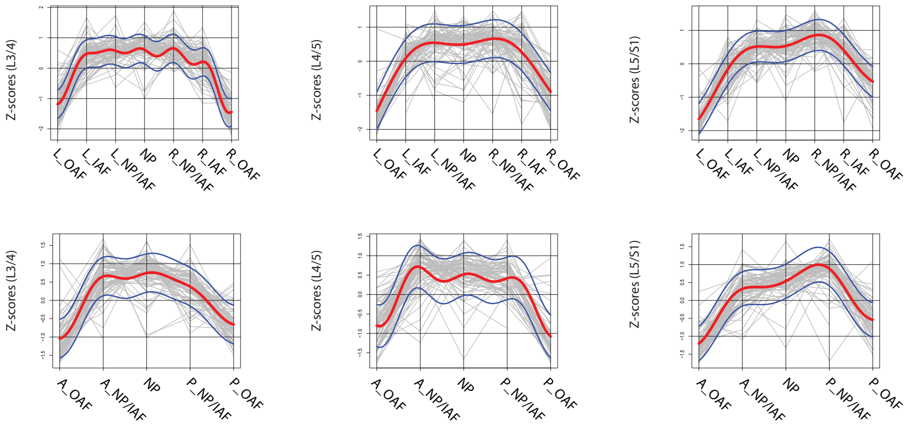


B

**Proteins in Y2 (n=171):**

HSP90B1, **CD81**, GDI2, PGLS, RAB1A, KRT2, KRT9, RPS21, SDCBP, GOT1, HNRNPC, F12, RAP1B, ARF4, ANXA4, ANXA6, DDOST, COMP, CILP, BGN, DSG1, OGN, PRELP, GNAI2, ANXA5, PKM, TFG, SRPX2, DCN, LUM, NES, ANXA7, H2AFV, ANXA1, GSTP1, CTSD, BPGM, YWHAQ, F11, SOD1, PRDX2, F13A1, CAT, IGHV5-51, SKP1, APEX1, **HIST1H1B**, CTSG, APOC3, TUBB, HBA1, CA2, SELENBP1, CA1, HBG1, TAGLN, ACTG1, CLIC1, HSP90AA1, SERPINB1, YWHAZ, CFL1, HIST1H4A, HBD, HBB, SERPINA7, SPARC, AK1, GPI, RPS27A, COL1A1, LAMA2, DPYSL3, IDH2, COL4A2, HSPB1, HUWE1, SH3BGRL3, ATP5A1, ATP5B, HIST2H2AC, AOC2, CILP2, DEFA3, GAPDH, MPZ, MSN, LOX, EIF4A1, APOL1, SERPINB6, RPL3, CAPZA2, UBA1, ASPN, MMP3, RPS8, ARF1, IGKV4-1, RPL18, **NT5E**, ACTN1, CAP1, CALM3, HSPA1B, HSPA8, C4A, YWHAE, SDHA, VWA1, COL6A2, COL6A1, COL6A3, FMOD, HNRNPA2B1, COL12A1, COL14A1, AHNAK, LMNA, AMBP, SERPINF1, LAMC1, ITIH1, LAMB2, NID1, THBS4, THBS1, THBS2, MYH9, PLS3, CALR, PFN1, SND1, HIST1H2BL, HNRNPA1, MYL6, MYL12A, SERPINH1, PDIA3, ANXA2, RPS3, NCL, PPIA, VCP, XRCC6, FSCN1, IQGAP1, CYB5R3, TAGLN2, PDIA6, FBLN1, MFGE8, TGFBI, CFD, MAMDC2, MATN2, LMNB2, LAMA4, SBSPON, DPYSL2, ACTN4, VAT1, FLNA, CDSN, SPTBN1, SPTAN1, TPM4, TUBA1B, VCL, ACO2, PAPPA

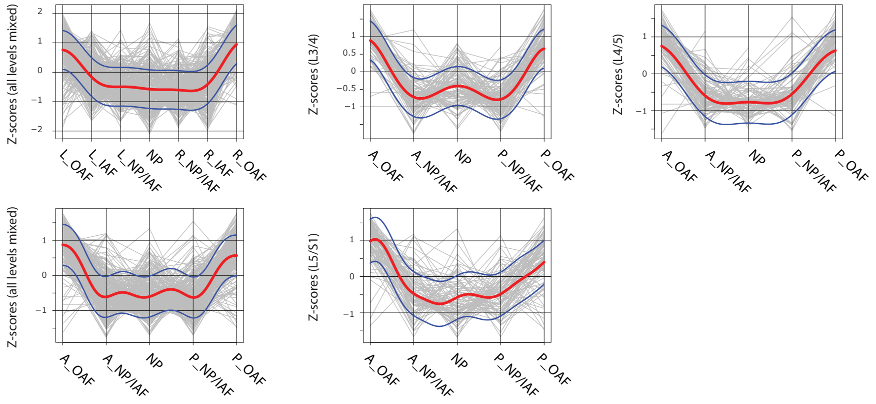


C

**Proteins in Y3 (n=25):**

ANGPTL7, RPN2, ALDOA, ENO3, CKM, CA3, MB, MYBPC1, MYL2, MYH2, TNNC1, ACTN2, MYH1, MYH7, MYLPF, MYL3, TPM2, TPM3, PYGM, TNNT3, TPM1, TPT1, ACLY, HHIP, NAP1L1

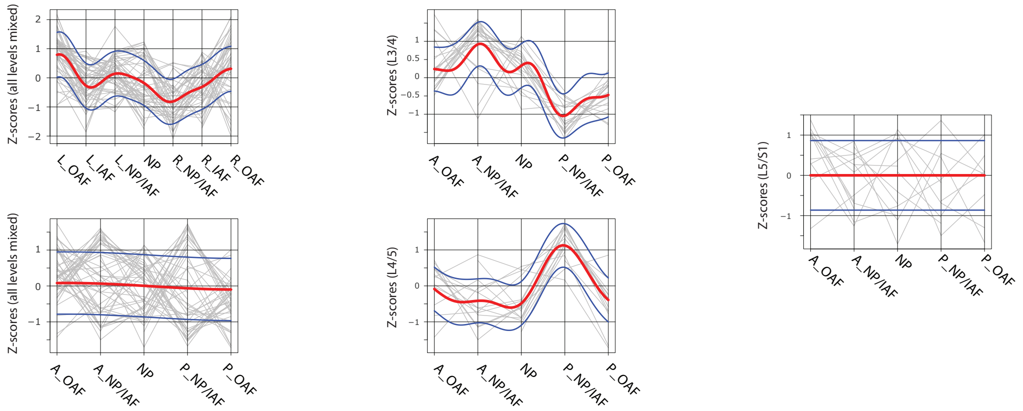


D

**Proteins in Y4 (n=48):**

RCN2, PGAM2, C8B, HP, APOA4, AKR1B1, C1QB, PGLYRP2, APOB, IGHM, SERPINF2, ITIH4, APCS, ITIH2, PLG, IGHG4, ITIH3, LBP, SERPINA4, C8G, C4BPA, PLIN4, IGHG3, IGLC6, C8A, GC, ORM1, C9, A1BG, HLA-DRB1, HSPD1, LRG1, F2, KNG1, ORM2, FGA, APOA1, FGB, FGG, IGHA1, HRG, HPX, TTR, VTN, IGHG1, ALB, IGHG2, TF

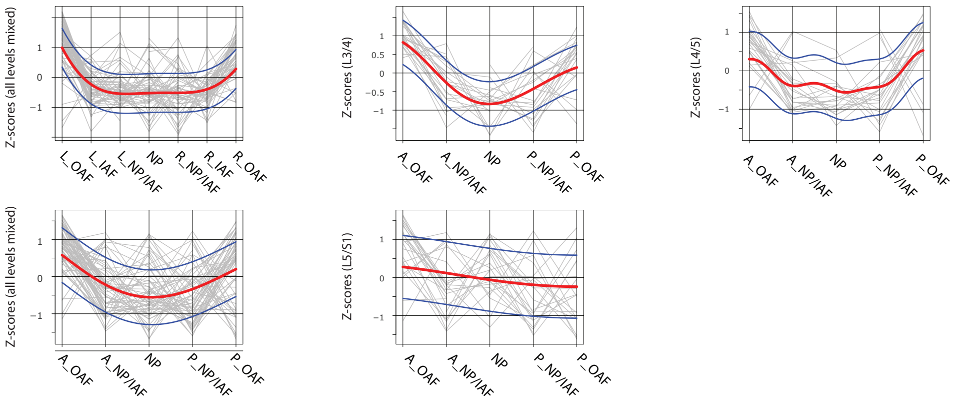


**Figure S8.** Proteins in modules Y1 (A), Y2 (B), Y3 (C) and Y4 (D), and their directional trends, in the young profiles. The red curve is the Gaussian Process Estimation (GPE) trend line, and the blue curves are 1 standard deviation above or below the trend line. Genes in red are transcription factors or DNA binding proteins. Genes in blue are surface markers.

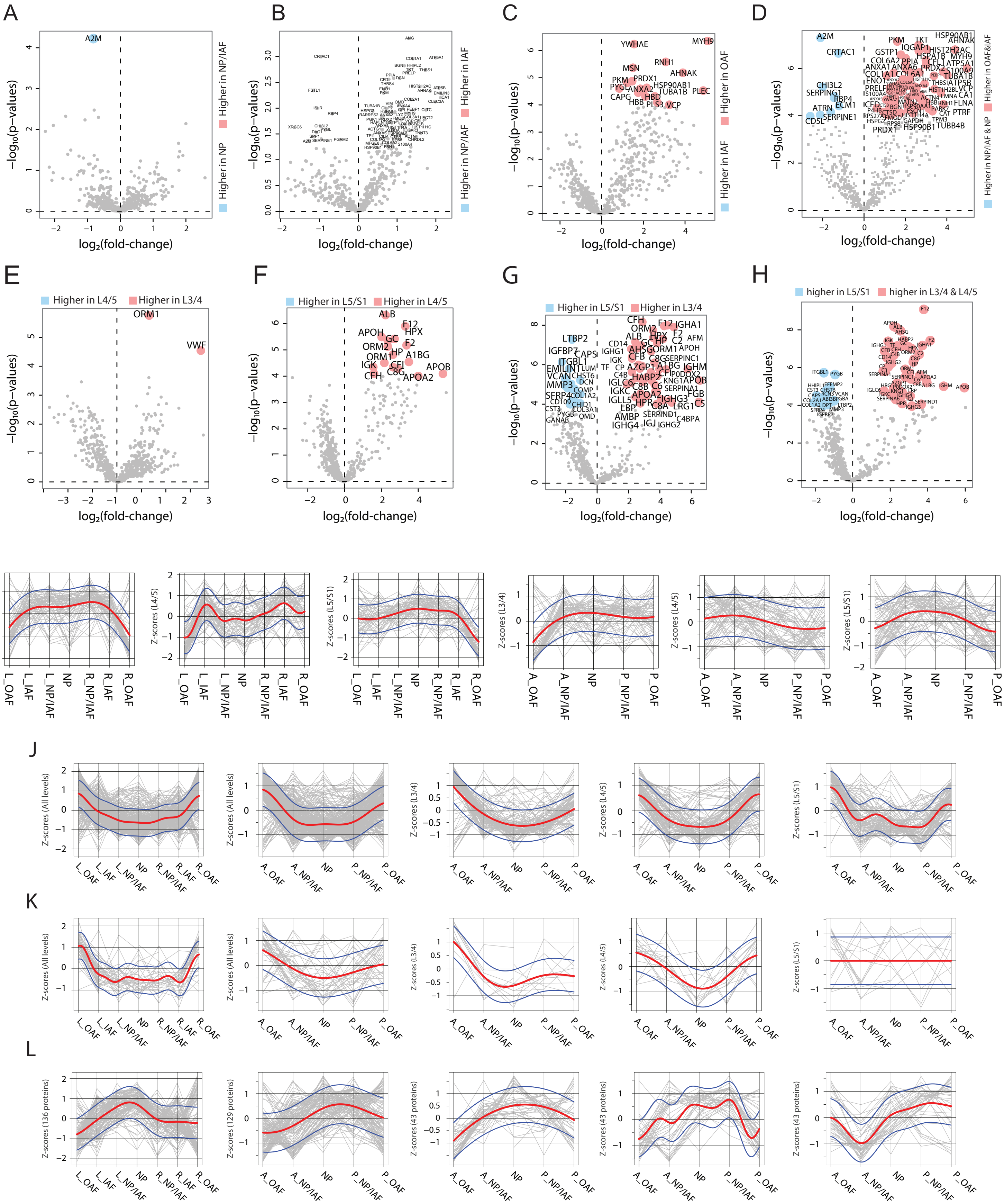


**Figure S9.** (A)-(H) Volcano plots showing the DEPs between different compartments or levels in the aged discs. (A) Volcano plot of DEPs between NP and NP/IAF. (B) Volcano plot of DEPs between NP/IAF and IAF. (C) Volcano plot of DEPs between IAF and OAF. (D) Volcano plot of DEPs between {NP + NP/IAF} and {IAF + OAF}. (E) Volcano plot of DEPs between L4/5 and L3/4. (F) Volcano plot of DEPs between L5/S1 and L4/5. (G) Volcano plot of DEPs between L5/S1 and L3/4. (H) Volcano plot of DEPs between lower level (L5/S1) and upper two levels combined, in the aged discs. (I)-(L), the lateral and anteroposterior trends of the four protein modules identified in (Figure S8) in the aged discs. (I) module Y1. (J) module Y2. (K) module Y3. (L) module Y4. The red curve is the Gaussian Process Estimation (GPE) trend line, and the blue curves are 1 standard deviation above or below the trend line.

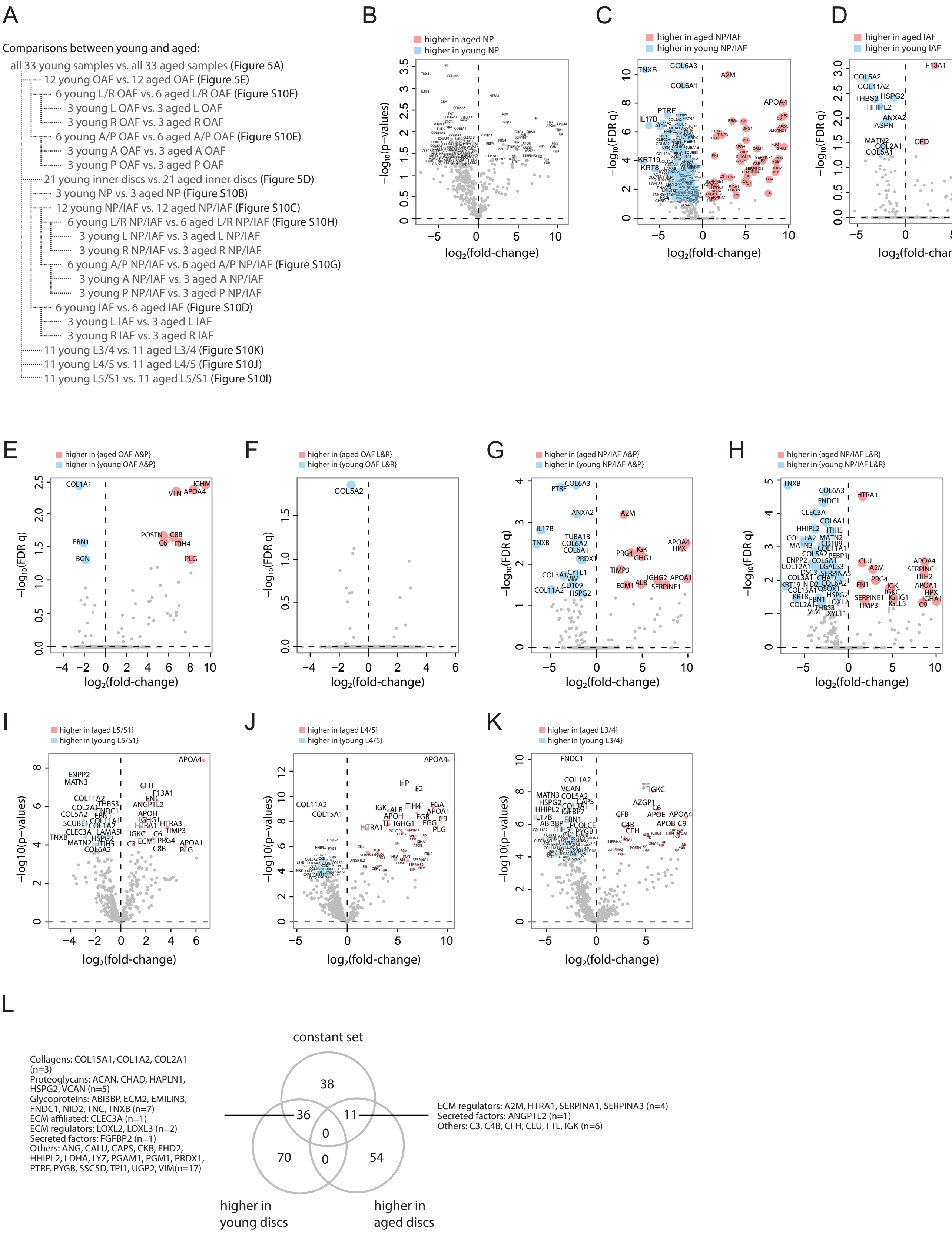


**Figure S10.** (A) Schematic diagrams showing the comparisons between young and aged profiles. (B)-(K) Volcano plots showing the differentially expressed proteins for each comparison listed in (A). (L) Venn diagram showing the overlaps of the DEPs between all young and all aged discs (from Figure 5A) and the constant set in the young discs (Figure 3D).

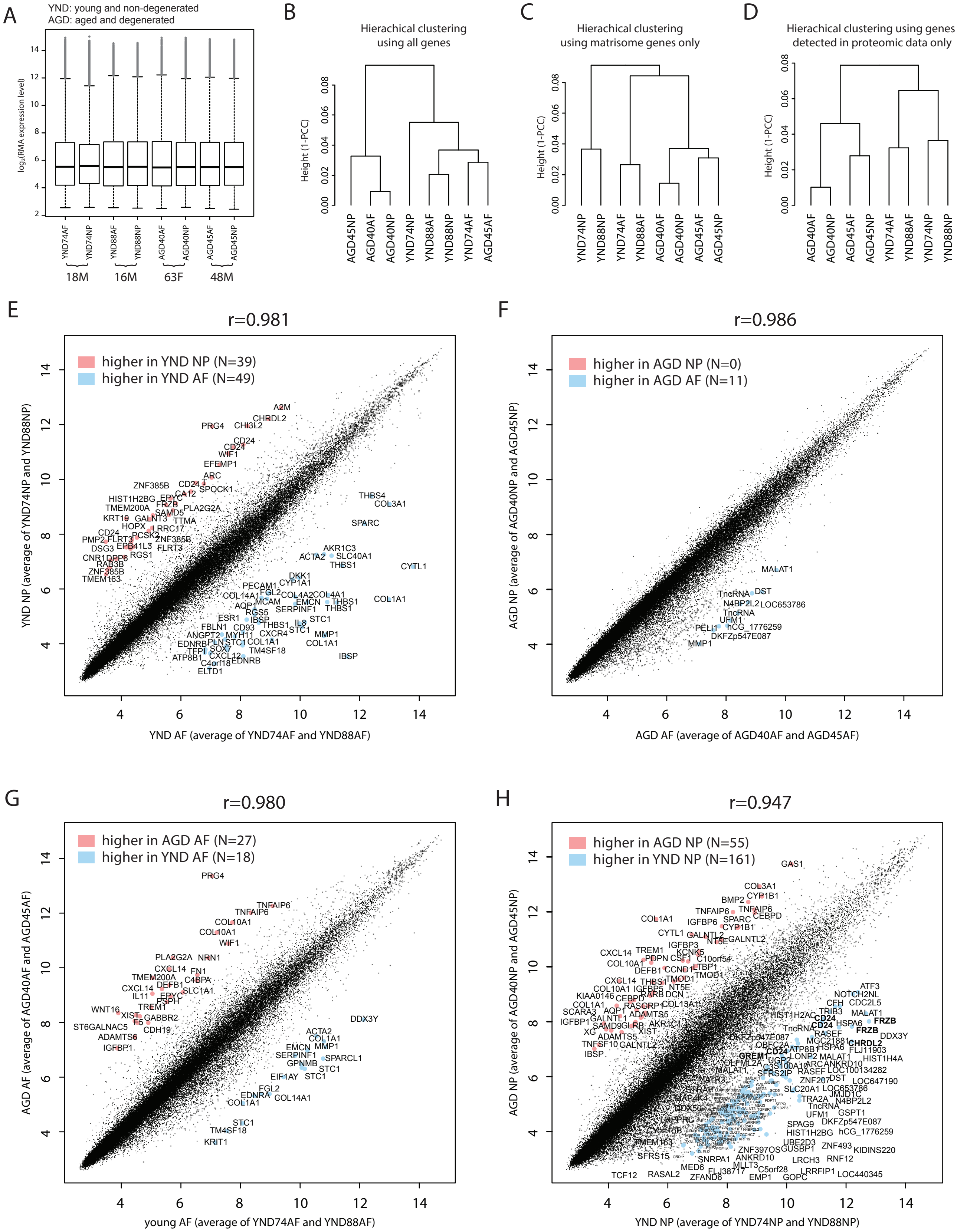


**Figure S11.** Microarray transcriptomic data of the NP and AF from four individuals, two of which are young and non-degenerated and the other two aged and degenerated. (A) boxplots of the normalized data show per-sample distribution of genome-wide expression profiles. (B)-(D) Hierachical clustering of the 8 microarray samples, based on genome-wide genes (B), matrisome genes (C), or the genes detected by proteomic data only (D). (E)-(H) Scatter plots of probesets between the average expressions of two groups. Red color indicates higher expression in the y-axis samples, and blue indicates higher expression in the x-axis samples. A $\log_2$(fold-change) of >3 and average expression >10 were used as cutoffs. Multiple instances of a differentially expressed gene are due to multiple probesets design of the array.

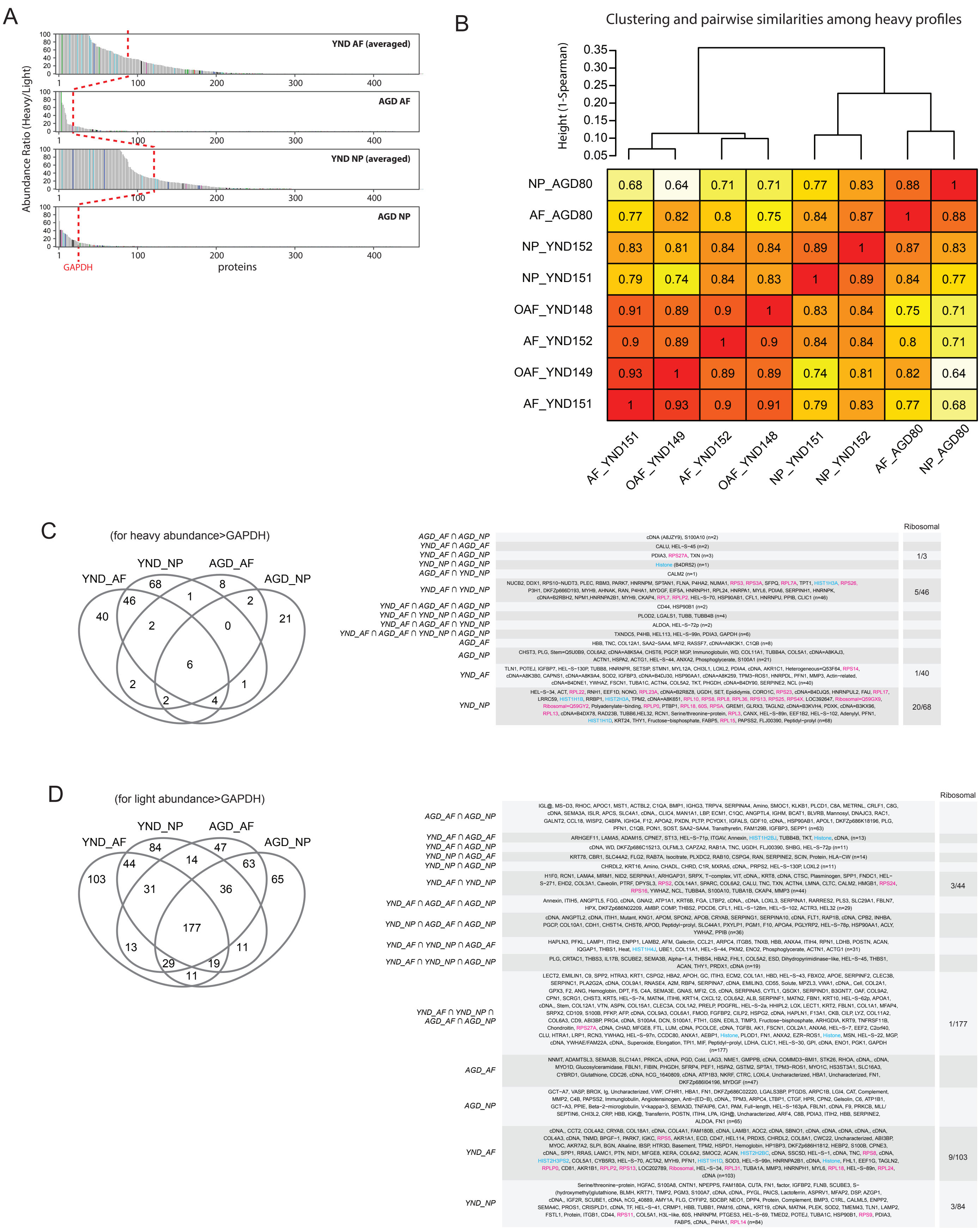


**Figure S12.** SILAC data. (A) The heavy-to-light ratios for each of the four groups were averaged, and then plotted in descending order of abundance. The red dotted reference line shows the expression of GAPDH. (B) Clustering (based on 1-Spearman as distance metrics and complete linkage) of the 8 heavy SILAC profiles shows that 'NP_YND151' and 'NP_YND152' have the tendency to cluster together, despite their difference in the numbers of detected proteins (Figure 7B left). Numbers in cells are Spearman correlation coefficients (based on non-missing values in both profiles under comparison) between pairs of profiles. (C) Venn diagram showing the overlap of proteins detected by the heavy SILAC profiles with abundance greater than GAPDH. The specific proteins in overlap are shown in the table to the right. (D) Venn diagram showing the overlap of proteins detected by the light SILAC profiles with abundance greater than GAPDH. The specific proteins in overlap are shown in the table to the right. Ribosomal proteins are highlighted in red.

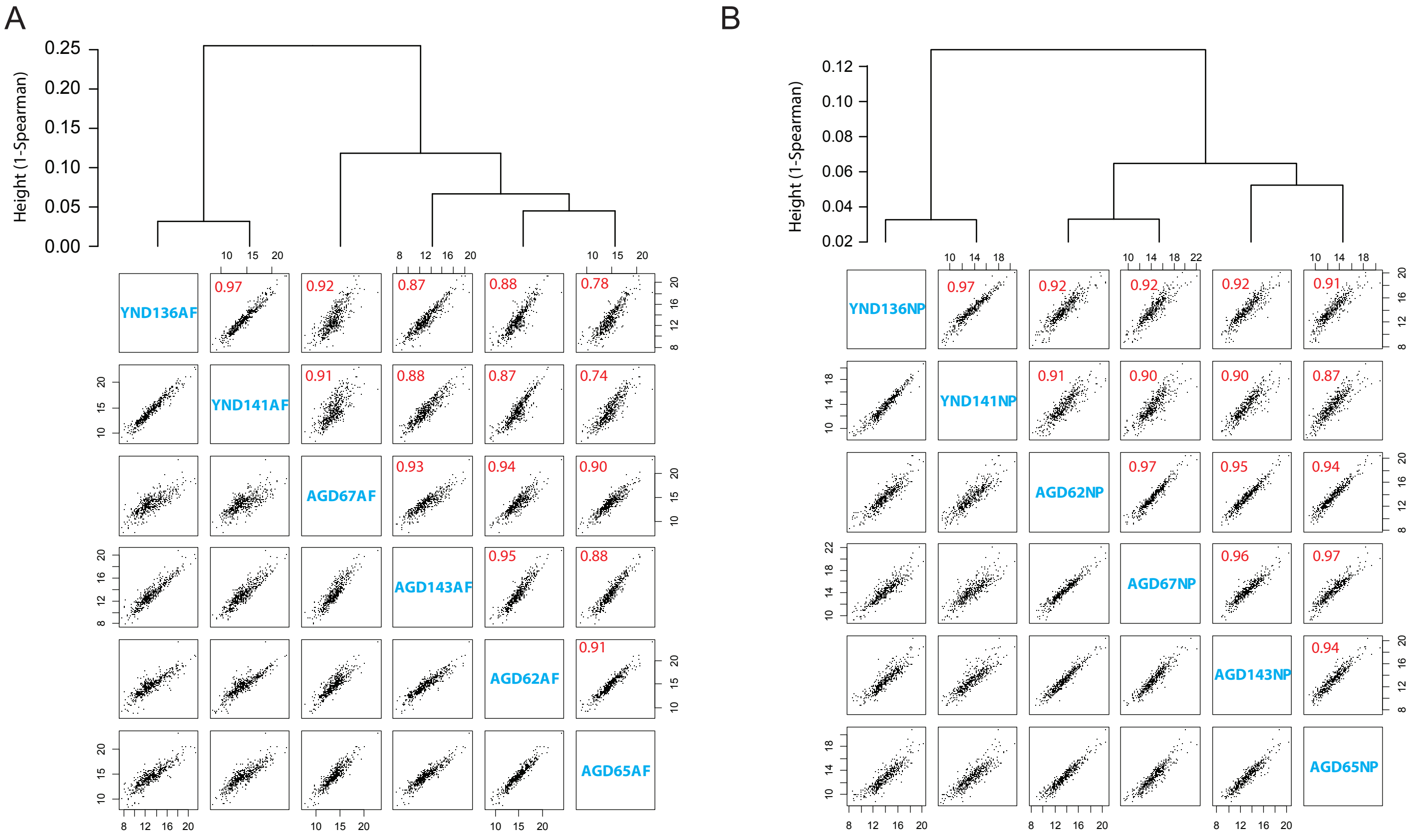


**Figure S13.** Degradome data. (A) Hierarchical cluster (upper panel) and pairwise scatter plot (lower panel) of the degradome profiles in the AF. Numbers in red are the Spearman correlation coefficient. (B) Hierarchical cluster (upper panel) and pairwise scatter plot (lower panel) of the degradome profiles in the NP. Numbers in red are the Spearman correlation coefficient.

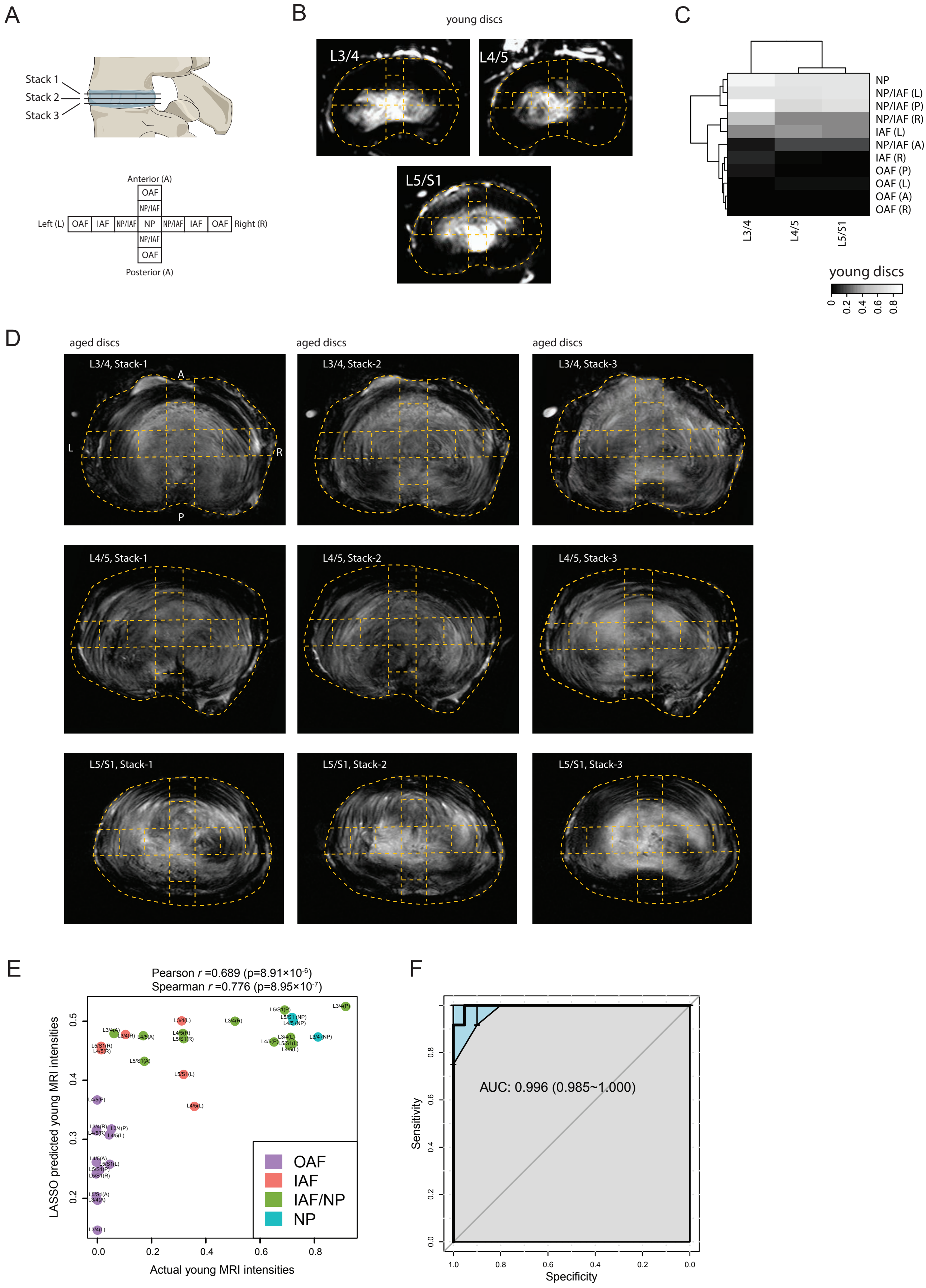


**Figure S14.** MRI-molecule connections. (A) Diagram showing the stacks of MRI per disc level and the 11 locations per disc. (B) Dashed curves overlaying the young discs' 3T MRI images, showing the compartments taken for proteomic profiling. (C) A heatmap with compartment and level bi-clustering, sshowing the relationship between regional MRI intensities. (D)Stacks of MRI images of the aged sample. (E) Scatter-plot showing the actual original MRI intensities of the young discs, and their predicted intensities of an LASSO model trained based on the ECM proteins most correlated with the aged disc MRI intensities. (F) A receiver operating characteristic (ROC) curve of the predicted MRI intensities between inner disc regions and OAF. AUC, area under the curve.
